# Autoimmune regulator act in synergism with thymocyte adhesion in the control of lncRNAs in medullary thymic epithelial cells

**DOI:** 10.1101/2021.07.14.452339

**Authors:** Max Jordan Duarte, Romário S. Mascarenhas, Amanda Freire Assis, Pedro Paranhos Tanaka, Cesar A. Speck-Hernandez, Geraldo Aleixo Passos

## Abstract

The autoimmune regulator (Aire) gene in medullary thymic epithelial cells (mTECs) encodes the AIRE protein, which interacts with its partners within the nucleus. This “Aire complex” induces stalled RNA Pol II on chromatin to proceed with transcription elongation of a large set of messenger RNAs and microRNAs. Considering that RNA Pol II also transcribes long noncoding RNAs (lncRNAs), we hypothesized that Aire might be implicated in the upstream control of this RNA species. To test this, we employed a loss-of-function approach in which Aire knockout mTECs were compared to Aire wild-type mTECs for lncRNA transcriptional profiling both *in vitro* and *in vivo* model systems. RNA sequencing enables the differential expression profiling of lncRNAs when these cells adhere *in vitro* to thymocytes or do not adhere to them as a way to test the effect of cell adhesion. Sets of lncRNAs that are unique and that are shared *in vitro* and *in vivo* were identified. Among these, we found the Aire-dependent lncRNAs as for example, Platr28, Ifi30, Morrbid, Malat1, and Xist. This finding represents the first evidence that Aire mediates the transcription of lncRNAs in mTECs. Microarray hybridizations enabled us to observe that temporal thymocyte adhesion modulates the expression levels of such lncRNAs as Morrbid, Xist, and Fbxl12o after 36h of adhesion. This finding shows the existence of a synergistic mechanism involving a link between thymocyte adhesion, Aire, and lncRNAs in mTECs that might be important for immune self-representation.

## 1. Introduction

The transcriptional control of gene expression in medullary thymic epithelial cells (mTECs) is a complex process, and according to recent findings, it involves two upstream controllers. One of these controllers is the autoimmune regulator (Aire) gene, which encodes the AIRE protein that functions as a transcriptional modulator, rather than as a classical transcription factor. Two features of the mode of action of AIRE classify it as a transcriptional modulator. First, AIRE associates with nuclear partner proteins, which interact with one another to form the AIRE complex. Next, the complex helps the stalled RNA Pol II to advance on chromatin and favors the transcription elongation phase in mTECs (Giraud et al., 2012), regulating a large set of downstream genes (Abramson et al., 2010; Passos et al., 2015, 2018; Perniola, 2018).

Forebrain embryonic zinc-finger-like protein 2 (Fezf2/FEZF2) is the second element that plays a role as a classical transcription factor that recognizes and interacts with promoter sequences on DNA (Takaba et al., 2015; Takaba and Takayanagi, 2017). The AIRE and FEZF2 proteins, although they display different modes of action, function in synergy, and both are under the control of the chromatin remodeler Chd4 in promoting transcriptional control at the chromatin level (Tomofuji et al., 2020).

In addition, the Aire gene is also involved in posttranscriptional control in mTECs by regulating the expression of miRNAs, which, in turn, negatively regulate downstream mRNAs (Macedo et al., 2013; Ucar et al., 2015; Oliveira et al., 2016).

Notable advances have been made in understanding and characterizing the mammalian cell transcriptome. As previously discussed (Assis et al., 2014), in functional terms, only a portion of the mammalian genome encodes RNAs, from which only a small fraction encode proteins; therefore, most RNAs are nonprotein-coding. Among the various species of mammalian RNAs described to date, a class of noncoding RNAs measuring more than 200 nucleotides, known as long noncoding RNAs (lncRNAs), have been attracting attention recently (Yao et al., 2019; Gil and Ulitsky, 2020; Bhatti et al., 2021a; 2021b; Song et al., 2021).

LncRNAs play roles as regulatory molecules that interact with other molecules, such as stretches of genomic DNA, proteins, miRNAs, or mRNAs (Zhang et al., 2018). The lncRNAs are implicated in such biological processes as normal eukaryotic development (Jarroux, Morillon and Pinskaya, 2017), cell differentiation and pluripotency (Chen et al., 2017; Yan et al., 2017), cancer biology (Bhan et al., 2017), the inflammatory response (Mathy and Chen, 2017) and autoimmunity (Zemmour, 2017). Among the subcategories of lncRNAs, intergenic lncRNAs, or simply lincRNAs, play a variety of roles, ranging from chromatin remodeling and RNA stabilization to the regulation of transcription (Ransohoff, Wei and Khavari, 2018).

However, the roles played by lncRNAs in the normal functioning of the thymus gland or in isolated thymic epithelial cells (TECs) have not been fully elucidated, and to the best of our knowledge, the few published results to date concern the effect of beta-estradiol on lncRNA expression in mTECs (Wei et al., 2018), thymic tumors (Su et al., 2020), effect of Zika virus on thymus (Messias et al., 2020) and thymic involution in birds (Li et al., 2021).

In this study, we hypothesized that Aire plays a role in the upstream control of lncRNAs in mTECs. To test this hypothesis, we used large-scale gene expression methods, such as next-generation RNA sequencing (RNA-Seq) and microarray hybridizations, to profile the lncRNA transcriptome of Aire wild type or Crispr-Cas9-generated Aire knockout (KO) mTECs in an *in vitro* model system.

Since the transcriptional activity of mTECs and their functionality is guaranteed through cell adhesion (Pezzi et al., 2016; Speck-Hernandez et al., 2018; Cotrim-Sousa et al., 2019; Irla, 2019), we also monitored the expression profiles of lncRNAs during the *in vitro* temporal adhesion to thymocytes.

Moreover, to evaluate the extension of the Aire effect on lncRNAs *in vivo*, we analyzed data of mTECs isolated of Aire wild type or of Aire KO mice.

## 2. Materials and Methods

### 2.1 Mice and separation of thymocytes

We employed C57BL/6 mice from the Jackson Laboratory (Bar Harbor, ME, USA) that were housed and mated at the Central Animal Facility, University of São Paulo, Campus of Ribeirão Preto, SP, Brazil, in temperature-controlled (approximately 22°C) and specific-pathogen-free conditions in 0.45-μm air-filtered ventilated racks under 12-h dark/light cycles and receiving water and food *ad libitum*.

For the surgical removal of the thymus and further separation of thymocytes, we employed four- to five-week-old females weighing 18-22 g using a previously described protocol (Donate et al., 2013). Before surgery, each animal was individually placed into an acrylic cage connected through a silicon pipe to a steel cylinder containing pressurized CO_2_, and the animal was euthanized by gas inhalation.

Thymi were removed from the animals, dissected and fragmented in RPMI 1640 medium. The thymocytes were collected by 2-3 passages of the thymic fragments through a 10-μm mesh nylon membrane (Sefar Inc., Depew, NY, USA) and centrifuged. The pelleted thymocytes were resuspended in phosphate-buffered saline (PBS) pH 7.4.

Fluorescence-activated cell sorting (FACS) analysis was performed using a BD FACSCalibur (BD Biosciences) flow cytometer with a phycoerythrin (PE)-labeled anti-CD3 antibody (Biolegend CNS, Inc. San Diego, CA, USA) and indicated that this procedure yielded a thymocyte population with a purity of ≥ 90%. These cells were utilized for further cell adhesion assays. All experiments with mice had been previously analyzed and approved by the Animal Research Ethics Committee, University of São Paulo, Campus of Ribeirão Preto, Brazil (approval number 003/2017-1).

### 2.2 Aire wild-type and knockout medullary thymic epithelial cell lines (mTECs)

For the *in vitro* experiments, we employed the murine (M. musculus) Aire wild-type (WT) mTEC line (EpCAM^+^, Ly51^−^, UEA1^+^) of mTEC 3.10 cells, as previously described (Nihei et al., 2003; Hirokawa et al., 1986). Besides it features the EpCAM^+^, Ly51^−^, UEA1^+^ phenotype, these cells also express Aire (Oliveira et al., 2013; 2016), Ccl21 and Sap1 genes (this work from the RNA-seq or microarray data) and the Aire protein (Supplemental material Figure 1). According to recent mTEC characterizations (Ribeiro et al., 2019), the mTEC 3.10 cell line is compatible with the mTEC^low^ differentiation stage.

Moreover, we previously have shown that the Aire WT mTEC 3.10 line express the Aire protein, which is localized in the nucleus of mTECs as shown through immunofluorescence (Pezzi et al., 2016; Speck-Hernandez et al., 2018).

In addition, we used a Crispr-Cas9-generated Aire knockout (KO), which was derived from the wild type mTEC 3.10 cell line and is known as the mTEC 3.10E6 clone.

The mTEC 3.10E6 clone was characterized as a KO compound heterozygous. This cell line is a carrier of indel mutations targeting exon 3 of the Aire gene and affecting both alleles, which feature different mutations. In Aire allele 1, there are two types of mutations: a T > G substitution (at position 351 of the encoded mRNA) followed by a nine-bp deletion (GCTGGTCCC, at positions 352-360 of the encoded mRNA) that results in the transcription of a 1,647-nucleotide Aire mRNA. In Aire allele 2, a single G deletion corresponds to position 352 of the encoded mRNA (resulting in a 1,655-nucleotide Aire mRNA). The T > G substitution in allele 1 causes the encoded AIRE protein to have a silent mutation (L118L).

In contrast, the nine-base-pair deletion caused a frameshift with a deletion of three amino acid residues (A119_P121del) and, consequently, a shorter and nonfunctional 548-amino-acid AIRE protein (Speck-Hernandez et al., 2018). FASTA sequences of Aire exon 3 are available at GenBank NCBI (https://www.ncbi.nlm.nih.gov/nucleotide/) under accession numbers MG493266 for Aire mutant allele 1 and MG493265 for Aire mutant allele 2. WT or KO mTECs were employed for further cell adhesion assays.

The Aire KO mTEC 3.10E6 mutant cell line does not localize Aire protein in the nucleus as previously shown through immunofluorescence (Speck-Hernandez et al., 2018).

For the *in vivo* model system analysis, we retrieve previously generated RNA-Seq data from mTECs (mTEC^high^ or mTEC^low^) isolated from Aire wild type or Aire KO mouse fresh thymi (St Pierre et al., 2015) whose raw data is available at Gene Expression Omnibus (GEO) available at (https://www.ncbi.nlm.nih.gov/geo/) under the accession number GSE65617.

### 2.3 mTEC-thymocyte cell adhesion assay

To evaluate the effect of deleterious Aire mutations (KO) in mTECs on thymocyte adhesion, we utilized a previously established protocol (Savino et al., 2004, 2010; Oliveira et al., 2016) with modifications, as described (Pezzi et al., 2016): specifically, the mTEC cells were cultured in RPMI 1640 medium containing 10% fetal bovine serum and antibiotics at 37°C in a 5% CO_2_ atmosphere. Semiconfluent cells were detached from their culture flasks by conventional trypsin/EDTA treatment, washed once with PBS at room temperature, resuspended in RPMI 1640 medium, and seeded in new culture flasks (2 × 10^6^ cells per 75 cm^2^ Corning® cell culture flasks). For this study, WT mTEC 3.10 cells or Aire KO mTEC 3.10E6 cells were cocultured with thymocytes, as described below.

Thymocytes were added to WT mTEC 3.10 or Aire KO mTEC 3.10E6 cultures at a ratio of 50:1 (thymocyte:mTEC) and cocultured in RPMI medium containing 10% fetal bovine serum and antibiotics at 37°C in a 5% CO_2_ atmosphere. In a typical adhesion experiment, the cell input was adjusted as follows: 5 × 10^6^ thymocytes and 1 × 10^4^ mTECs. This gives a 50: 1 ratio (thymocytes: mTECs). Next, the nonadherent thymocytes were carefully removed from cultures by washing with PBS at 37°C and discarded. The culture flasks were washed more vigorously with PBS at 4°C to remove the adherent thymocytes, which were kept for counting and phenotyping. Single-positive (SP) CD4^+^ or CD8^+^ cells were stained with an APC-Cy7-labeled anti-CD4 and FITC-labeled anti-CD8 antibody (Biolegend), respectively, and quantified using a BD LRSFortessa flow cytometer (BD Biosciences).

Next, mTEC cells were detached from their culture flasks by conventional trypsin/EDTA treatment and resuspended in PBS for counting. Cell counts for either thymocytes or mTECs were performed on a Cellometer Auto T4 Cell Viability Counter (Nexcelon Bioscience, Lawrence, MA, USA). Cells were recovered from cocultures after 12 or 36 h, which correspond to two incubation time points that give linear response in terms of thymocyte adhesion (data not shown).

Experiments were performed in triplicate (i.e., three co-cultures with Aire WT mTEC 3.10 and three cocultures with Aire KO mTEC 3.10E6) and repeated at least three times, with similar results being obtained. The adhesion index (AI) was calculated as follows: AI = number of adhered thymocytes/number of mTEC cells. Statistical analysis was performed by Student’s t-test using the GraphPad Prism 6.0 platform. P-values < 0.05 were considered to be significant.

### 2.4 Total RNA preparation and reverse transcription real-time quantitative PCR (RT-qPCR)

We followed previously described protocols (Pezzi et al., 2016; Cotrim-Sousa et al., 2019). Briefly, total RNA was prepared using the mirVana® kit (Ambion, Austin, TX, USA) according to the manufacturer’s instructions. Through UV spectrophotometry, RNA preparations were confirmed to be free of proteins and phenol. RNA degradation was assessed by microfluidic electrophoresis using Agilent RNA 6000 nanochips and an Agilent 2100 Bioanalyzer (Agilent Technologies, Santa Clara, CA, USA). Only RNA samples that were free of proteins and phenol and had RNA integrity numbers (RINs) ≥ 9.0 were employed for cDNA synthesis with SuperScript reverse transcriptase enzyme according to the manufacturer’s instructions (Invitrogen Corporation, Carlsbad, CA, USA).

RT-qPCR was employed to confirm the expression levels of Fendrr, Peg13, Fam219aos, Platr28 and Neat1 lncRNAs. The expression level of each lncRNA was normalized to the housekeeping mRNA Hprt; these gene is commonly used as reference. The Primer3 web tool (http://frodo.wi.mit.edu/primer3) was utilized to select pairs of oligonucleotide primers spanning an intron/exon junction with an optimal melting temperature of 60°C.

The cDNA sequences of lncRNAs were retrieved from the NCBI GenBank Database (http://ncbi.nlm.nih.gov/nuccore?itool=toolbar). The forward (F) and reverse (R) sequences (presented in the 5’ to 3’ orientation) were the following: mRNA Hprt (NM_013556.2) CCCCAAAATGGTTAAGGTT and CAAGGGCATATCCAACAACA, lncRNA Fendrr (NR_130109) CTCTCCAGTTCCCACCACC and TGGTCTGGTTCTGGTCACTC, lncRNA Peg13 (NR_002864)TGTGACCACGAACCGAAGAG and TCGTCTACATAGCACCAGCG, lncRNA Fam219aos (NR_045726) AGGCAGAGGTTCTGATTCACAG and GTGAAAAACTGACCCCCTGGA, lncRNA Platr28 (NM_001018010) TTGGAGCACAAAATGGGACCT and GGGCCAAGCTCAGGTCAAC, lncRNA Neat1 (NR_003513) CTGCACTGTAGATCGGGACC and TCCCCAACACCCACAAGTTT.

Transcription was quantified using a StepOne Real-Time PCR System (Applied Biosystems, USA). The analyses were performed using the cycle threshold (Ct) method, which enables quantitative analysis of the expression of a factor using the formula 2-DΔC^t, in which ΔCt = Ct target cDNA sequence - Ct of the housekeeping mRNA Hprt, and ΔΔCt = ΔCt sample - ΔCt.

### 2.5 Western blot of AIRE protein

Western blotting (WB) of AIRE protein expressed in Aire wild type mTEC 3.10 or Aire KO mTEC 3.10E6 cell lines, was performed according a previously published protocol optimized in our laboratory (Pezzi et al., 2016). Briefly, we use a primary anti-AIRE goat polyclonal antibody (Anti-AIRE D17 IgG antibody, Santa Cruz Biotechnology Inc., Dallas, TX, USA) and a horseradish peroxidase (HRP)-conjugated donkey anti-goat IgG as secondary antibody (Merck Millipore, Darmstadt, Germany).

The WB polyvinylidene fluoride (PVDF) membrane (BioRad, Hercules, CA, USA) was developed using a peroxidase substrate for chemiluminescence (LuminataForte™, Merck Millipore) in an ImageQuant™ LAS 500 apparatus (GE Life Sciences, Piscataway, NJ, USA).

Protein bands were visualized using the ImageQuant™ GE Life Sciences (Piscataway, NJ, USA) and quantitatively analyzed with ImageJ software (version 1.49). AIRE protein was normalized to GAPDH protein, which is routinely used as a housekeeping. The same membrane that was used to detect AIRE protein was washes and reused in an incubation with an anti-GAPDH rabbit polyclonal primary antibody (Cell Signaling Technology, Beverly, MA, USA) following incubation with peroxidase conjugated anti-rabbit antibody.

The membrane was developed as described above. Statistical comparison of protein bands intensity between in Aire wild type mTEC 3.10 and Aire KO mTEC 3.10E6 cell lines was performed using Student’s t-test within the GraphPadPrism 6.0 platform (http://www.graphpad.com/prism/Prism.htm).

### 2.6 RNA sequencing (RNA-Seq)

This analysis was based on the protocol as previously described (Speck-Hernandez et al., 2018). Briefly, paired-end (2 × 100 bp) sequencing was performed using an Illumina HiSeq 2500 sequencer (Illumina, San Diego, CA, USA) installed at LacTad Laboratory Facility, Campinas State University, Campinas, SP, Brazil. To this end, we employed a TruSeq Stranded Total RNA Library Prep Kit (Illumina).

Data were analyzed following this process: the quality of raw FASTQ sequences was first analyzed through a FASTQC program (https://www.bioinformatics.babraham.ac.uk/projects/fastqc/). FASTQC sequences were mapped to the Mus musculus reference genome (mm10) using the STAR 2.5 Spliced Aligner program (https://github.com/alexdobin/STAR), which output a BAM file containing the sequences and their genomic references, as well as a GFT file with gene annotations employed for further determinations of the number of reads per transcript through the HTSeq Count program (https://htseq.readthedocs.io/en/master/).

The edgeR package (https://bioconductor.org/packages/release/bioc/html/edgeR.html) was employed to analyze the differentially expressed (DE) lncRNAs in Aire KO mTEC 3.10E6 compared to Aire WT mTEC 3.10 cells before and after thymocyte adhesion. We considered DE lncRNAs with fold changes ≥ 2.0 and adjusted P-values (FDR) ≤ 0.05. The RNA-Seq data obtained in this work are available on GEO under the accession number GSE91015.

For the analysis of DE lncRNAs expressed *in vivo* by mTECs isolated of Aire wild type or Aire KO fresh thymi, we used the same pipeline as above described.

### 2.7 Microarray hybridization and data analysis

We utilized the protocol established by our group and previously published (Donate et al., 2013; Oliveira et al., 2013; Fornari et al., 2015). Briefly, the gene expression changes were evaluated using the Agilent one-color (Cy3 fluorochrome) microarray-based gene expression platform according to the manufacturer’s instructions. To hybridize mRNAs or lncRNAs to whole-genome 8 × 60 K 60-mer oligonucleotide arrays (Agilent Technologies, Palo Alto, CA, USA), 200 ng of total RNA was labeled with the one-color Quick Amp labeling kit (Agilent Technologies).

Samples of complementary RNA (cRNA) were hybridized for 18 h at 42°C in a rotator oven and washed. The array slides were scanned using a DNA microarray scanner (Agilent Technologies), and the hybridization signal was extracted using Agilent Feature Extraction software (version 10.5). Gene expression profiles from two or three independent preparations of control (mTEC 3.10 cells) and Aire KO (mTEC 3.10E6 cells) before or after thymocyte adhesion were analyzed by comparing the microarray hybridizations of the respective samples.

A complete file, including all of the oligo sequences present in the microarray representing mRNAs or lncRNAs employed in this study and the experimental conditions, is available online in the ArrayExpress public database (https://www.ebi.ac.uk/arrayexpress/) under accession number E-MTAB-6541.

The quantitative microarray data were normalized to the 75th quantile and analyzed using the R platform (https://www.r-project.org/) (version 3.10). Computational tools contained in the arrayQualityMetrics (https://www.bioconductor.org/packages/release/bioc/html/arrayQualityMetrics.html) (Kauffmann, Gentleman and Huber, 2009), and Agi 4 × 44 PreProcess (López-Romero et al., 2010), (http://www.bioconductor.org/packages//2.11/bioc/manuals/Agi4x44PreProcess/man/Agi4x44PreProcess.pdf) were utilized for the preprocessing of the data. Bioconductor packages contain algorithms for assessing the quality of the data, background correction, and normalization. After analyzing the DE mRNAs or lncRNAs, functions contained in the limma package (https://bioconductor.org/packages/release/bioc/html/limma.html) (Wettenhall and Smyth, 2004), were employed.

This study considered only those mRNAs or lncRNAs with adjusted P-values ≤ 0.05, controlled false discovery rates (FDR) by the Benjamini-Hochberg method, and fold changes ≥ 1.5 to be DE. The DE lncRNAs were hierarchically clustered. Heat maps were constructed using Euclidean distance and the complete linkage method to evaluate controls and Aire KO mTECs’ expression profiling before and after thymocyte adhesion. As described above for RNA-seq data, we utilized the lncTAR (http://www.cuilab.cn/lnctar) and CPC2 (http://cpc2.cbi.pku.edu.cn/) for the functional enrichment of lncRNAs.

### 2.8 Identification and nomenclature of differentially expressed lncRNAs

The identification of lncRNAs was performed according to their annotations in the UCSC Genome Browser (https://genome.ucsc.edu/) and Agilent’s annotation regarding the microarray slides.

### 2.9 Identification and genomic positioning of protein-encoding genes near lncRNAs

The R platform (https://www.r-project.org/) and biomaRt package (https://www.ensembl.org/biomart/martview/) were utilized to access databases, including Ensembl (https://www.ensembl.org/), to search in the genome the protein-coding genes closest to the lncRNAs.

### 2.10 Evaluation of the coding potential of lncRNAs

The coding potential of differentially expressed lncRNAs was analyzed using CPC2 software (http://cpc2.cbi.pku.edu.cn/), a calculator that assesses the coding potential of a given lncRNA using an alignment-free technique. This software employs a logistic regression model based on four characteristics: 1) the size of the open reading frame (ORF, or open reading matrix), since long ORFs are rarely observed in noncoding sequences; 2) the ORF coverage, that is, the ratio between ORF and transcript lengths; 3) the Fickett TESTCODE score, which reflects the possible combinatorial effect of the nucleotide composition and the tendency to use a particular codon; and 4) the hexamer score, which determines the relative degree of bias in the use of a hexamer in a given sequence (Kong et al., 2007; Kang et al., 2017).

The values of this score vary from 0 to 1. Values close to 0 indicate lower coding potential, and as they approach 1, they indicate more significant coding potential based on the cutoff of 0.5 (Kang et al., 2017). We analyzed the coding potential only of the differentially expressed lncRNAs cataloged at Ensembl (https://www.ensembl.org/index.html).

## 3. Results

### 3.1 mTEC-thymocyte adhesion is influenced by Aire

For the cell adhesion assay, we used murine (*M. musculus*) mTECs (Aire WT mTEC 3.10 or Aire KO mTEC 3.10E6 cell lines).The mTEC-thymocyte coculture for 12 or 36 h showed a significant reduction in the number of adhered thymocytes on Aire KO mTEC 3.10E6 cells as demonstrated throughout a conventional cell adhesion assay and whose mTECs and thymocytes were visualized by light microscopy. In order to calculate the adhesion index (AI) both cell types from cocultures were individually counted. Since in physiological conditions and within the thymus, single positive (SP) thymocytes adhere to mTECs, we use flow cytometry analysis to show that *in vitro* an equal proportion between SP CD4^+^ and CD8^+^ thymocytes were adhered to this cell type. Thymocyte adhesion was lowered with Aire KO mTEC 3.10E6 and was not preferred for one of the SP subtypes (Figure 1).

**Figure 1.**
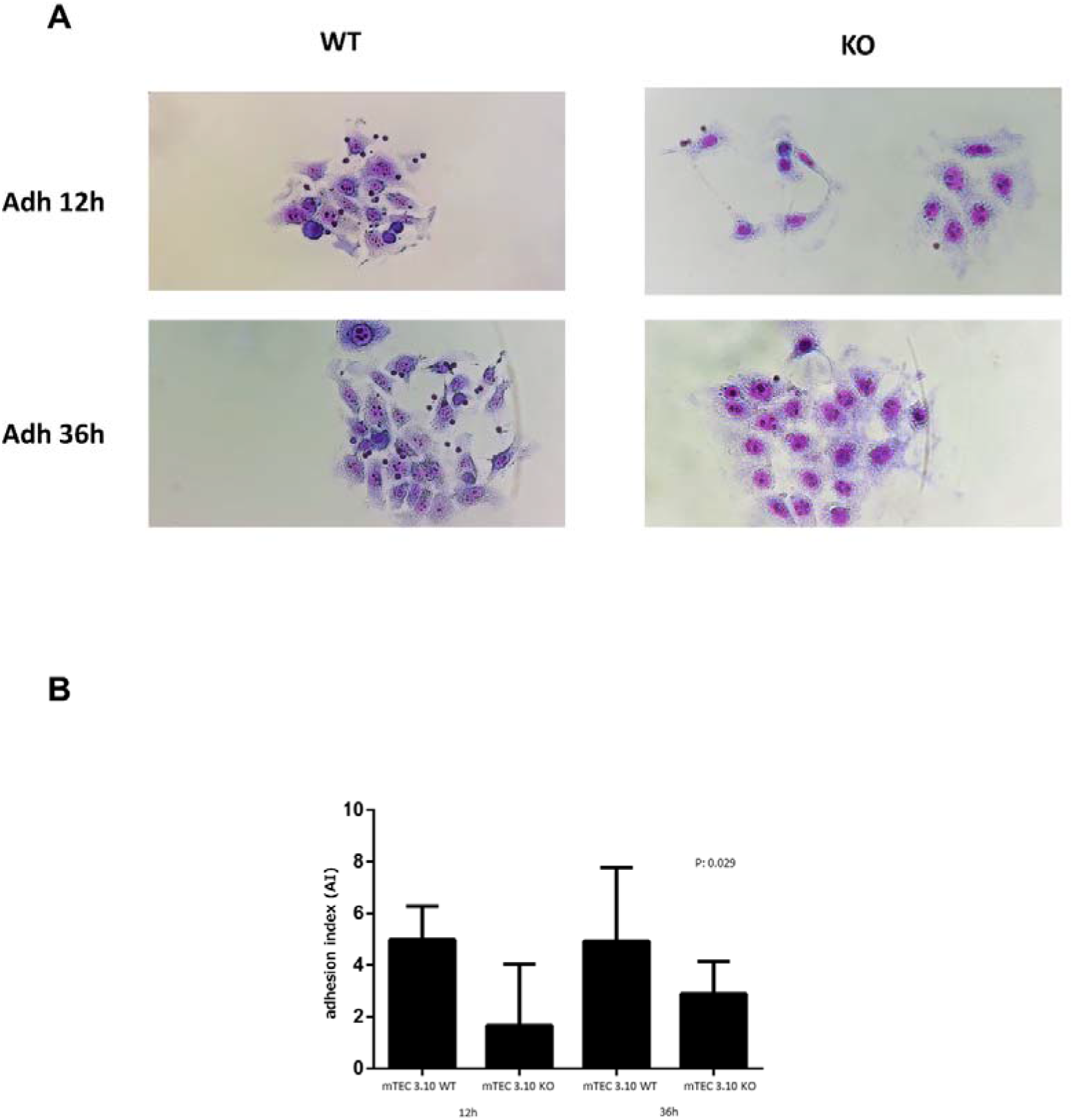
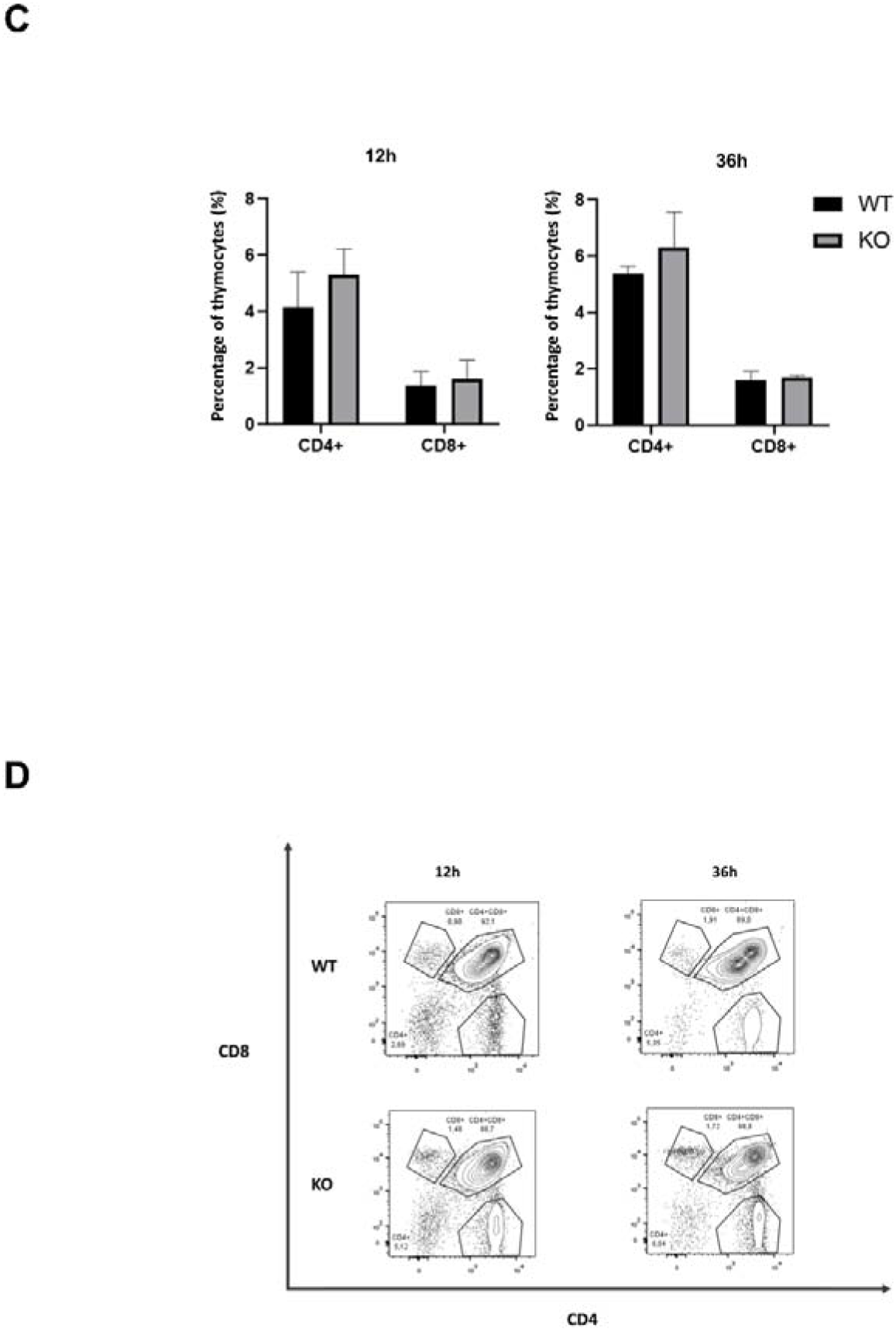
The mTEC-thymocyte adhesion was comparing Aire wild-type (WT) mTEC 3.10 with Aire KO mTEC 3.10E6. **(A)** Photomicrography of fresh mTEC-thymocyte co-cultures after 12h or 36h adhesion. Conventional light microscopy, Giemsa staining, 20 × magnification. **(B)** Adhesion indexes after 12 or 36 h mTEC-thymocyte co-cultures. **(C)** Quantification of CD4 and CD8 cell surface markers of adhered or non-adhered thymocytes to mTECs. **(D)** Flow cytometry of CD4 single-positive (SP) thymocytes and CD8 SP adhered or non-adhered thymocytes to mTECs and CD4/CD8 double-positive thymocytes adhered to Aire wild type (WT) mTEC 3.10.

### 3.2 Differential expression of lncRNAs is influenced by Aire and thymocyte adhesion

Initially we use RNA-Seq to identify the differentially expressed (DE) lncRNAs between mTEC samples. The 136 DE lncRNAs as observed in the unsupervised hierarchical clustering heat map of Figure 2, were considered Aire-dependent, of which 72 were upregulated, and 64 were downregulated when comparing Aire wild type (WT) vs Aire knockout (KO) mTECs subjected, or not, to thymocyte adhesion. This result demonstrate the influence of Aire and thymocyte adhesion on lncRNAs in mTEC cells. The top ten DE lncRNAs are presented in Supplemental material Table 1 and, Table 1.1.

**Figure 2.**
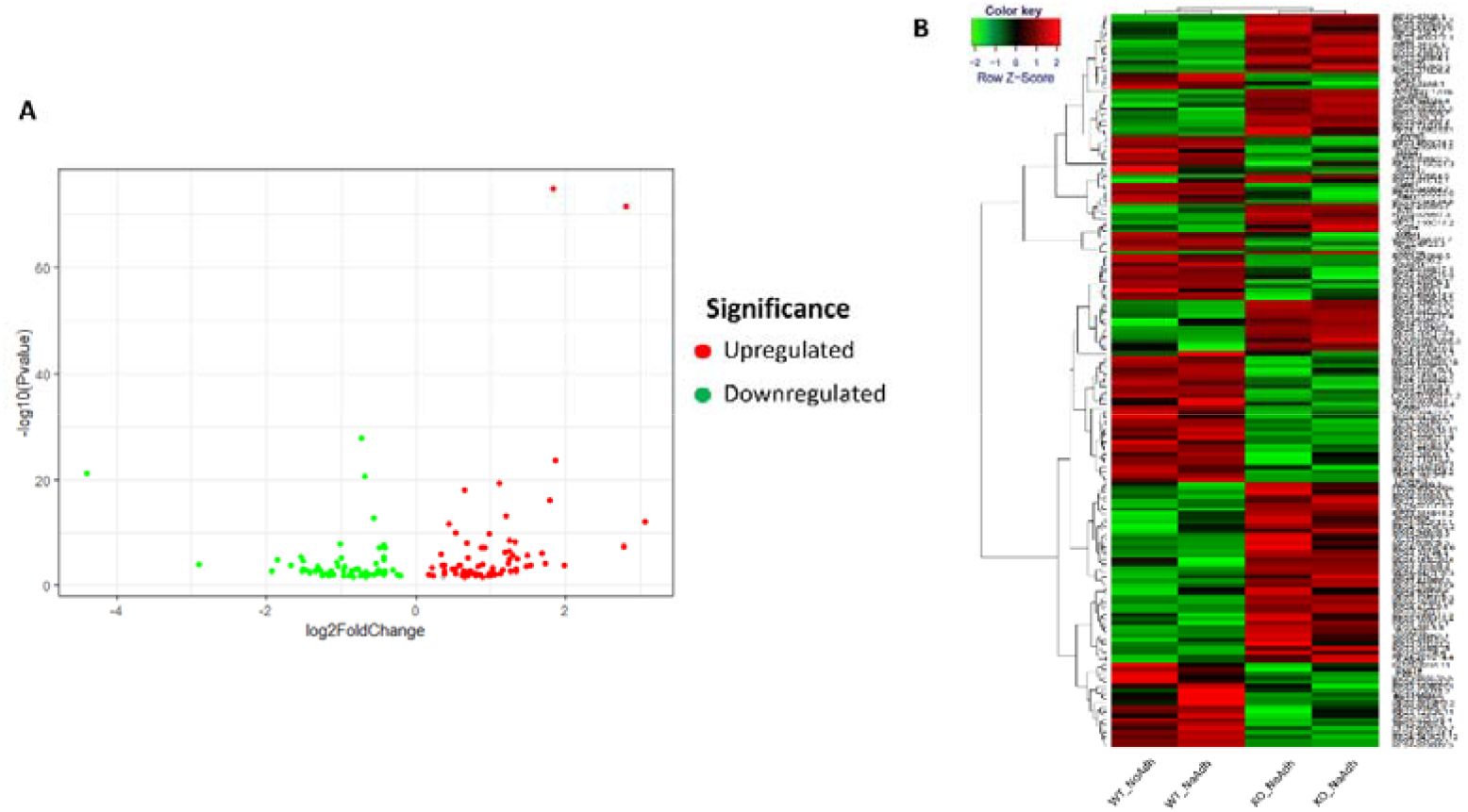
Differentially expressed lncRNAs were considered Aire-dependent, of which 72 were upregulated, and 64 were downregulated when comparing Aire wild type (WT) vs Aire knockout (KO) mTECs subjected, or not, to thymocyte adhesion. Total RNA samples were extracted from WT or Aire KO mTECs and sequenced through Illumina protocol (RNA-Seq). **(A)** Volcano plot showing the differentially expressed lncRNAs in Aire KO mTEC 3.10E6. Red dots represent the upregulated and green dots, the downregulated lncRNAs, fold-change ≥ 2.0. **(B)** Heat map showing differentially expressed lncRNAs in Aire KO mTEC 3.10E6. Red means the upregulated and green the downregulated lncRNAs. Row Z scores, fold-change ≥ 2.0 normalize heat-map colors.

#### 3.2.1 Effect of temporal thymocyte adhesion on mTEC lncRNA expression

In order to evaluate whether temporal thymocyte adhesion may cause differences in the expression profile of lncRNAs, we analyzed their expression throughout microarray hybridizations varying the coculture time. The Venn diagram depicted in Figure 3, shows 45 identified DE lncRNAs when comparing Aire WT mTEC 3.10 vs. Aire KO mTEC 3.10E6 cells before thymocyte adhesion. Of these, eight lncRNAs were upregulated, and 37 were downregulated. After 12h of thymocyte adhesion, 53 DE lncRNAs (16 up- and 37 downregulated) were observed, and after 36h of adhesion, 138 DE lncRNAs (31 up- and 107 downregulated) were identified (Figure 3).

**Figure 3.**
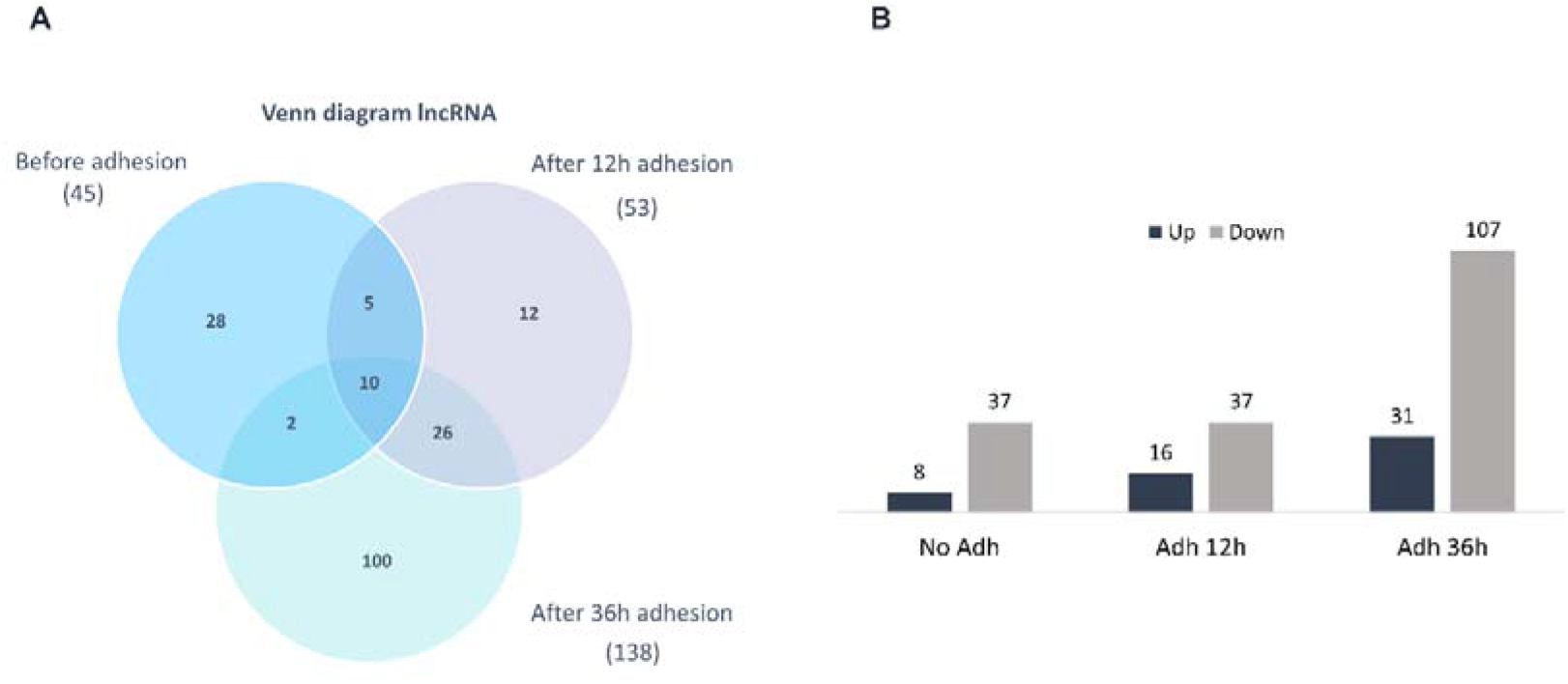
**(A)** Venn diagram of the differently expressed lncRNAs comparing Aire wild-type (WT) mTEC 3.10 with Aire KO mTEC 3.10E6 before and after thymocyte adhesion. **(B)** Summary of differently expressed lncRNAs. Total RNA samples were extracted from WT or Aire KO mTECs subjected or not to thymocyte adhesion during 12 or 36 h, labeled and hybridized to Agilent microarrays.

Figure 4 shows the hierarchical clustering of the transcriptional expression of lncRNAs, comparing Aire WT mTEC 3.10 with Aire KO mTEC 3.10E6 cells before and after temporal thymocyte adhesion, with adjusted P-values ≤ 0.05 (Benjamini-Hochberg FDR) and fold-changes ≥ 1.5 serving as the cutoff, as shown in the heat map.

**Figure 4.**
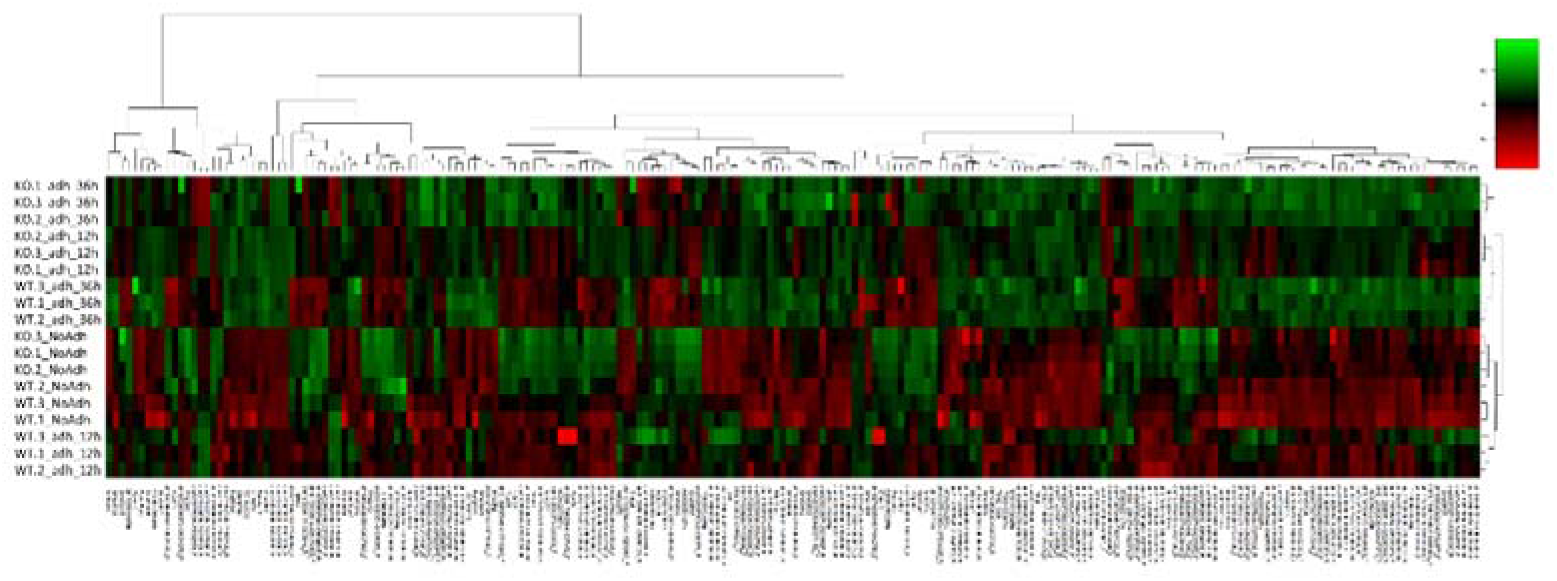
Heat map plot of differentially expressed lncRNAs comparing Aire wild-type (WT) mTEC 3.10 with Aire KO mTEC 3.10E6 before and after thymocyte adhesion. Total RNA samples were extracted from WT or Aire KO mTECs subjected or not to thymocyte adhesion during 12 or 36 h, labeled and hybridized to Agilent microarrays. Unsupervised dendrograms and heat-map were constructed using the R platform. Red = upregulated mRNAs, green = downregulated mRNAs and black = unmodulated mRNAs (Fold-change ≥ 2.0, false discovery rate FDR ≤ 0.05, Pearson correlation metrics, 75 percentile).

### 3.3 Extent of the effect of Aire in the lncRNA expression *in vitro* and in vivo

Then we asked if Aire could influence the expression of mTEC lncRNAs both *in vitro* and *in vivo*. For this, we compared the expression data of lncRNAs collected in this study that uses the *in vitro* mTEC-thymocyte adhesion with the expression of lncRNAs from mTECs separated from fresh mouse thymus. The Venn diagram depicted in Figure 5 shows that either *in vivo* or *in vitro* the absence of Aire influences the number of DE lncRNAs. As the expression data from Aire wild type mTECs were used as a reference, the Venn diagram was constructed with data from Aire KO mTECs in which appear the shared and the exclusive lncRNAs according to the sample analyzed, i.e., Aire KO mTEC^low^ vs Aire KO mTEC^high^ isolated from *in vivo* fresh thymus (diagram left side) or Aire KO mTEC without thymocyte adhesion vs Aire KO mTEC under *in vitro* thymocyte adhesion (diagram right side).

**Figure 5.**
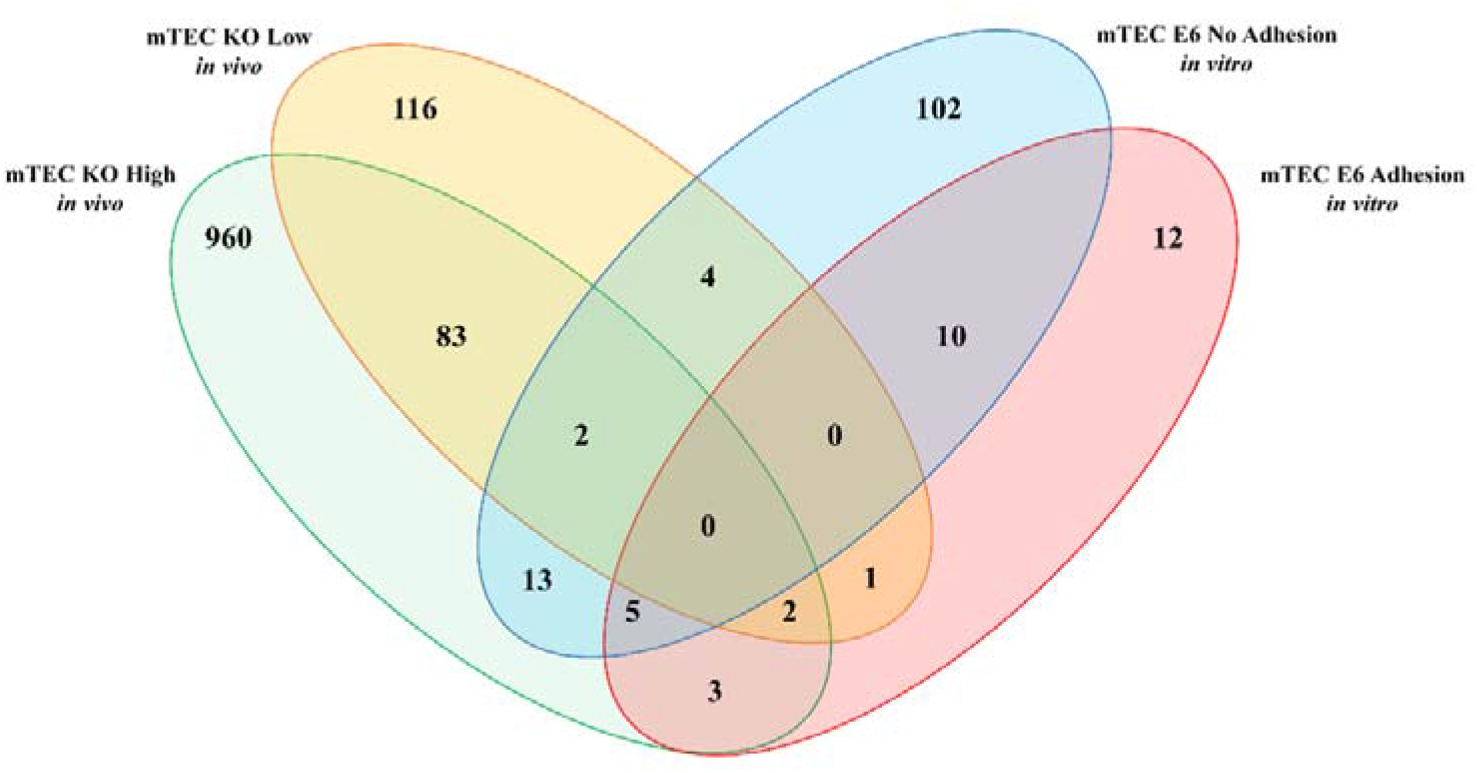
Venn diagram showing the exclusive and shared differentially expressed lncRNAs in Aire KO mTECs. *In vivo* Aire KO mTEC^low^ / Aire KO mTEC^high^ (left side) and *in vitro* Aire KO mTEC with no thymocyte adhesion / Aire KO mTEC under thymocyte adhesion (right side). The RNA-Seq data from the *in vivo* mTECs were retrieved from St Pierre et al., (2015) and from the *in vitro* mTECs, this study.

### 3.4 Positioning of protein coding genes near lncRNA genes

Regarding the relative position in the genome of protein-encoding genes to lncRNAs, we located a wide range both upstream and downstream considering the different chromosomes. For example, while the Neat1 lncRNA is found 29,796 bp upstream of the Frmd8 protein-encoding gene, this lncRNA is 74,059 bp downstream of the Scyl1x gene. Also, the lncRNA Meg3 is 59,329 bp upstream of the Rtl1 gene and 88,178 bp downstream of the Dlk1 gene (Supplemental material Table 2, Table 2.1 and, Table 2.2).

### 3.5 Coding potential of lncRNAs

Among the RNAs that are differentially expressed (DE) in mTECs before their adhesion with thymocytes, we found 13 that were unlikely to encode a protein and were therefore classified as lncRNAs (Supplemental material Table 3). After 12 h of thymocyte adhesion, we listed 27 differentially expressed RNAs, 26 of which were classified as lncRNAs along with an exception of the DNAH10-202. Although DNAH10-202 falls into this classification, it presents coding potential (Supplemental material Table 3.1). Next, after 36 h of thymocyte adhesion, we listed 56 differentially expressed lncRNAs, among which the following exceptions presented coding potential (1700018B24Rik, DNAH10-202 shared with 12-h adhesion, E230016K23Rik, Gm10069, Gm14636, Gm16062, Gm20324, and Gm50039) (Supplemental material Table 3.2).

### 3.6 Validation of the transcriptional expression levels by RT-qPCR

To confirm the modulation of selected lncRNAs we evaluated the modulation of Fendrr and Peg13 lncRNAs that were identified by RNA-Seq and Fam2019aos, Platr28, and Neat1 lncRNAs that were identified through microarray hybridizations (Supplemental material Figure 2).

## 4. Discussion

The autoimmune regulator (AIRE) protein complex, which is formed in the nucleus by the interactions between AIRE and its partner proteins (Abramson et al., 2010; Kyewski and Peterson, 2010), exerts its function by advancing stalled RNA Pol II on chromatin, thereby enabling the transcription elongation phase in which both coding and noncoding RNAs are synthesized (Giraud et al., 2012; Waterfield et al., 2014; Passos et al., 2018). This property guided us when we observed that the Aire gene could control miRNAs in medullary thymic epithelial cells (mTECs) (Macedo et al., 2013).

A possible association of Aire with posttranscriptional control in mTECs began to be considered in our group when we discussed the results obtained by the Giraud’s group (Giraud et al., 2012). There are findings indicating that AIRE protein pushes stalled RNA Pol II on chromatin. Since RNA Pol II transcribes miRNAs in addition to mRNAs, Aire might even indirectly control the expression of miRNAs (Macedo et al., 2013).

Given this possibility, we hypothesized that Aire could control lncRNAs in mTECs. As the link between mTECs, Aire and lncRNAs has not been explored to date, we realize the uniqueness of this proposition. Therefore, we established a model system that makes it possible to test this hypothesis. The primary mTEC 3.10 cell line of continuous growth in culture (Hirokawa et al., 1986; Mizuochi et al., 1992) and its Aire KO subclone (Aire^−/−^ mTEC 3.10E6) (Speck-Hernandez et al., 2018).

The *in vitro* adhesion with thymocytes significantly modulates the transcriptional activity of mTECs, upregulating the transcriptional expression of Aire and Fezf2, as well as cell adhesion-related genes such as Cd80 or Tcf7, among others (Speck-Hernandez et al., 2018; Cotrim-Sousa et al., 2019). Therefore, we ponder that the mTEC-thymocyte coculture comprises a model system, which is helpful for large-scale transcriptional gene expression studies influenced or not influenced by adhesion with thymocytes.

Considering that adhesion with thymocytes is an essential property of mTECs, we evaluated this variable because it also controls transcriptional gene expression. Compared to Aire wild-type mTECs, the Aire KO mTECs adhere less to the thymocytes of CD4^+^ or CD8^+^ phenotypes, which corroborated our previous findings (Speck-Hernandez et al., 2018) and validated the model system for this study.

The prepared RNA-Seq library enabled us to evaluate the lncRNA sequences that are differentially expressed when comparing Aire wild-type vs. Aire KO mTECs before or after thymocyte adhesion. This finding represents the first observation that Aire can control lncRNAs in mTECs with synergistic adhesion action with thymocytes. Therefore, the lncRNAs influenced by Aire are referred to in this report as Aire-dependent. Among the set of differentially expressed (DE) lncRNAs, also were found the sub-class of intergenic lncRNAs (lincRNAs) (Ransohoff et al., 2018). The lncRNA Ifi30 is Aire-dependent and is of particular interest, since it participates in antigen processing and presentation pathways (https://www.genome.jp/kegg/pathway.html); it might contribute to mTECs during self-antigen presentation.

It was possible to determine the chromosomal location of the differentially expressed lncRNAs and their closest coding genes. The genomic location is essential to predict the mode of action of lncRNAs; in other words, the lncRNAs can either act in cis, regulating genes close to their locus on the same chromosome, or in *trans*, when they regulate genes located in distant regions on different chromosomes (Gil and Ulitsky, 2020; Yan et al., 2017). We found that the lncRNAs Fendrr, Gas5, Malat1, Neat1, and Xist might, in the model system studied, act in cis, since their closest protein-encoding targets are located on the same DNA segment on chromosome either upstream or downstream (Bao et al., 2019).

The expression levels of the lncRNAs Fendrr, and Peg13, as measured by RNA-Seq, were further validated using RT-qPCR. The results obtained in this experiment corroborated the bioinformatic analysis of the RNA-seq dataset, confirming the quality and precision of the *in silico* analyses. Fendrr is one of the most well-characterized lncRNA at present (Grote, 2014). It has been shown that Fendrr acts in the development of the heart of mice and that its downregulation leads to heart failure (Grote, 2014). In human gastric cancer and renal carcinoma, Fendrr upregulation has been reported to be associated with a worse prognosis, as it promotes increased migration, proliferation, and invasion of cancer cells (He et al., 2019).

Therefore, this study is the first that associates the lncRNA Fendrr with the immune system, i.e., thymic stromal cells. It is worth pursuing this lncRNA to determine if, in the mTEC cells, it is involved in proliferation and/or migration processes that are essential in the biology of the thymus.

In previous studies performed in our laboratory, it was observed that both silencing or knocking out the Aire gene in mTEC 3.10 cells resulted in a decrease in the ability of these cells to adhere to thymocytes (Pezzi et al., 2016; Speck-Hernandez et al., 2018). Genes involved in cell-cell adhesion, such as Cd80, Itgam, Tgfbi, Cdh16 and Itgae had their expression levels reduced when Aire was disrupted.

Considering that the physical process of interaction between thymocytes and mTECs is crucial in presenting PTAs and the maturation of mTECs and T-cells themselves, the absence of Aire alters the entire connection structure, compromising the functioning of central immune tolerance (Derbinski et al., 2005; Passos et al., 2018; Perniola, 2018).

One crucial point is that mTEC-thymocyte crosstalk can influence the transcriptional profile of mRNAs as previously reported (Speck-Hernandez et al., 2018; Cotrim-Sousa et al., 2019) and in this study we hypothesized that lncRNAs might be modulated in such condition. Through the adhesion assay, it was possible to observe that in the culture of Aire KO mTEC 3.10E6 cells with thymocytes, varying the coculture time between 12 or 36 h, there was a decrease in adhesion compared with WT mTEC 3.10 cells. We found no cell morphology changes in visual observation and under an optical microscope between WT mTEC 3.10 cells and Aire KO mTEC 3.10E6 cells.

We were able in demonstrating that thymocyte adhesion also modulates lncRNAs in mTECs. After coculturing Aire WT mTEC 3.10 cells with thymocytes, we identified 191 differentially expressed lncRNAs, 53 after 12 h and 138 after 36 h of coculture. Among these RNAs, we focused on the lncRNA NEAT1, which plays a role in controlling the formation of paraspeckle heterochromatin structures (Clemson et al., 2009; Zhang et al., 2013). This result suggests that longer thymocyte adhesion stimulates lncRNA-mediated posttranscriptional control in mTECs.

During T-cell development, immature T-cells interact with the cell matrix of the thymus, which is composed primarily of type I and IV collagen, integrins, laminin, and fibronectin (Savino et al., 2004). Integrins are involved in the interaction with the extracellular matrix, physical cell-cell contact, and adhesion receptors. Integrins are essential in the processes that modulate the proliferation and differentiation of T-cells (Savino et al., 2004, 2013; Bertoni et al., 2018).

Interestingly, the deregulation of transcriptional expression of lncRNAs was observed in Aire KO mTECs, suggesting the involvement of Aire-lncRNAs axis with the expression of molecules linked to cell-cell and cell-matrix interactions.

Although not explored in this study, there may be posttranscriptional interaction between mRNAs that encode cell adhesion proteins and lncRNAs identified here.

To compare the extent to which lncRNAs are controlled by Aire in an *in vivo* system, we reanalyzed raw RNA-seq data previously collected from mTECs (mTEC^high^ and mTEC^low^) from Aire WT and Aire KO mice (St Pierre et al., 2015). It was possible to observe that both in an *in vitro* and *in vivo* model systems, there was modulation of lncRNAs using the approach of loss-of-function (LOF) of Aire (Aire KO). The mTEC subtypes that most expressed exclusive lncRNAs were Aire KO mTEC^high^ (*in vivo*) and Aire KO mTEC 3.10E6 (in vitro without adhesion with thymocytes). In addition, subsets of exclusive and shared lncRNAs were also observed between Aire KO mTEC^low^ (*in vivo*), Aire KO mTEC^high^ (*in vivo*), Aire KO mTEC 3.10E6 (*in vitro* without adhesion) and Aire KO mTEC 3.10E6 (*in vitro* under adhesion with thymocytes).

In conclusion, the results of this study demonstrate for the first time the existence of a link between Aire and the expression lncRNAs in mTEC cells. The adhesion between mTECs and thymocytes was shown to be synergistic in this process. Through the knockout of Aire, we were able to employ a LOF strategy and observe that downstream lncRNAs are deregulated.

Considering the regulatory role of lncRNAs, the axis Aire-lncRNAs in mTECs found in this study, might be important for the *in vivo* induction of central immunological tolerance.

## Data availability

High-quality paired-end reads of this study are available at the Gene Expression Omnibus (GEO) database under accession number GSE91015. The microarray data are available at the ArrayExpress database under accession number E-MTAB-6541.

## Acknowledgements

We thank Dr. Wilson Savino (Laboratory on Thymus Research, Oswaldo Cruz Institute, Fiocruz, Rio de Janeiro, Brazil) that gently ceded the Aire WT mTEC 3.10 line, Dr. Daniella A. Mendes-da-Cruz and Dr. Vinícius Cotta-Almeida (Laboratory on Thymus Research, Oswaldo Cruz Institute, Fiocruz, Rio de Janeiro, Brazil) and Dr. Eduardo A. Donadi (Ribeirão Preto Medical School, USP, Ribeirão Preto, Brazil) for help and discussions and MSc Denise B. Ferraz (Ribeirão Preto Medical School, USP, Ribeirão Preto, Brazil) for technical assistence.

This study was funded by São Paulo Research Foundation (Fapesp, São Paulo, Brazil, through grants # 13/17481-1 and # 17/10780-4 to GAP), Conselho Nacional de Desenvolvimento Científico e Tecnológico (CNPq, Brasília, Brazil, through grant # 305787/2017-9 to GAP). This study was financed in part by the Coordenação de Aperfeiçoamento de Pessoal de Nível Superior (CAPES, Brasília, Brazil) through Financial Code 001.

---

**Supplemental material Figure 1.**
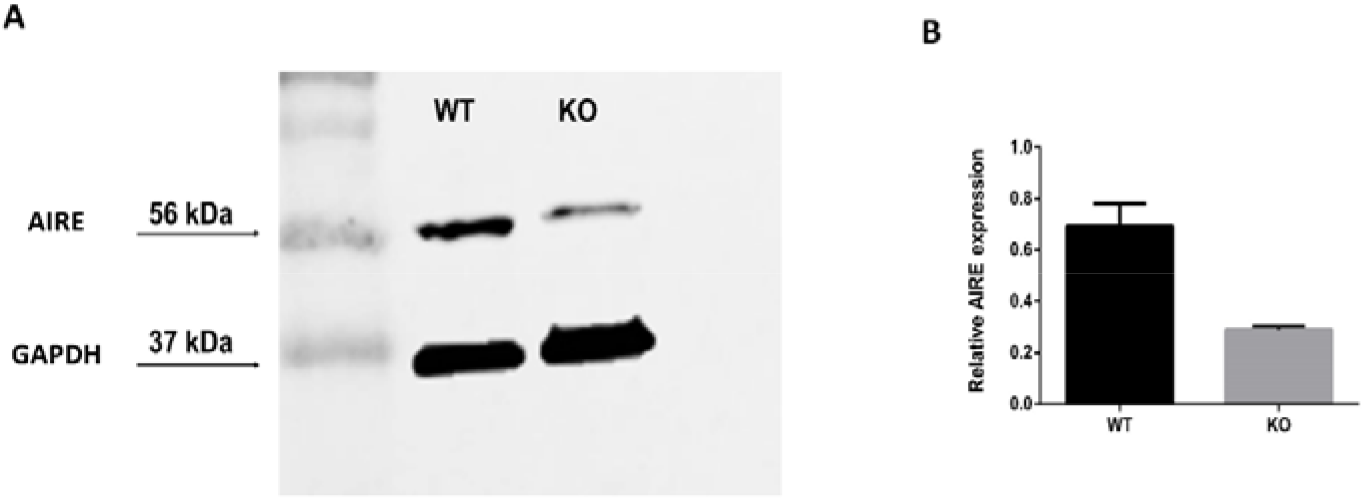
**(A)** Representative western blot after SDS-PAGE of Aire WT mTEC 3.10 and Aire KO mTEC 3.10E6 cell lysates, probed with an antibody against AIRE protein. GAPDH protein was used as an internal load control. **(B)** Western blot bands were quantified by ImageJ software, and AIRE protein levels (normalized to GAPDH) were expressed as signal intensity (pixels/area). The bar graph shows that the defective mutant AIRE protein is expressed at low levels in the Aire KO mTEC 3.10E6 cell line. The cell lysates were prepared from Aire WT mTEC 3.10 or Aire KO mTEC 3.10E6 cell lines with no adhesion with thymocytes.

**Supplemental material Figure 2.**
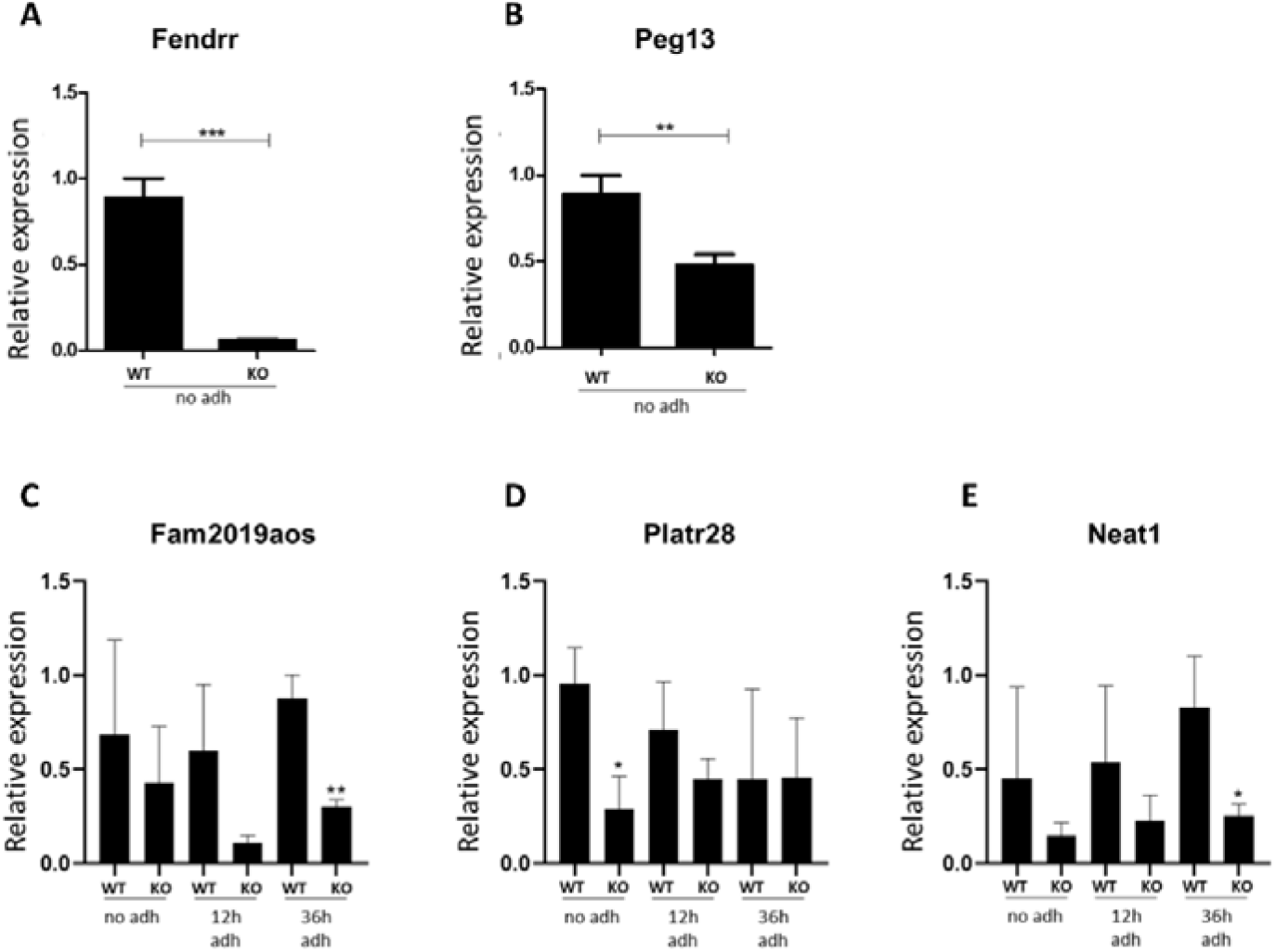
Validation of the relative transcriptional expression levels of the Fendrr, Peg13, Fam219aos, Platr28, Neat1 lncRNAs **(A-E)**, comparing Aire wild-type (WT) mTEC 3.10 with Aire KO mTEC 3.10E6 before and after thymocyte adhesion. The transcriptional expression of lncRNAs were quantified by real-time quantitative-PCR (RT-qPCR). Means ± S.D. from experiments from three independent replicates. *P* < 0.05.

**Supplementary Table 1.**
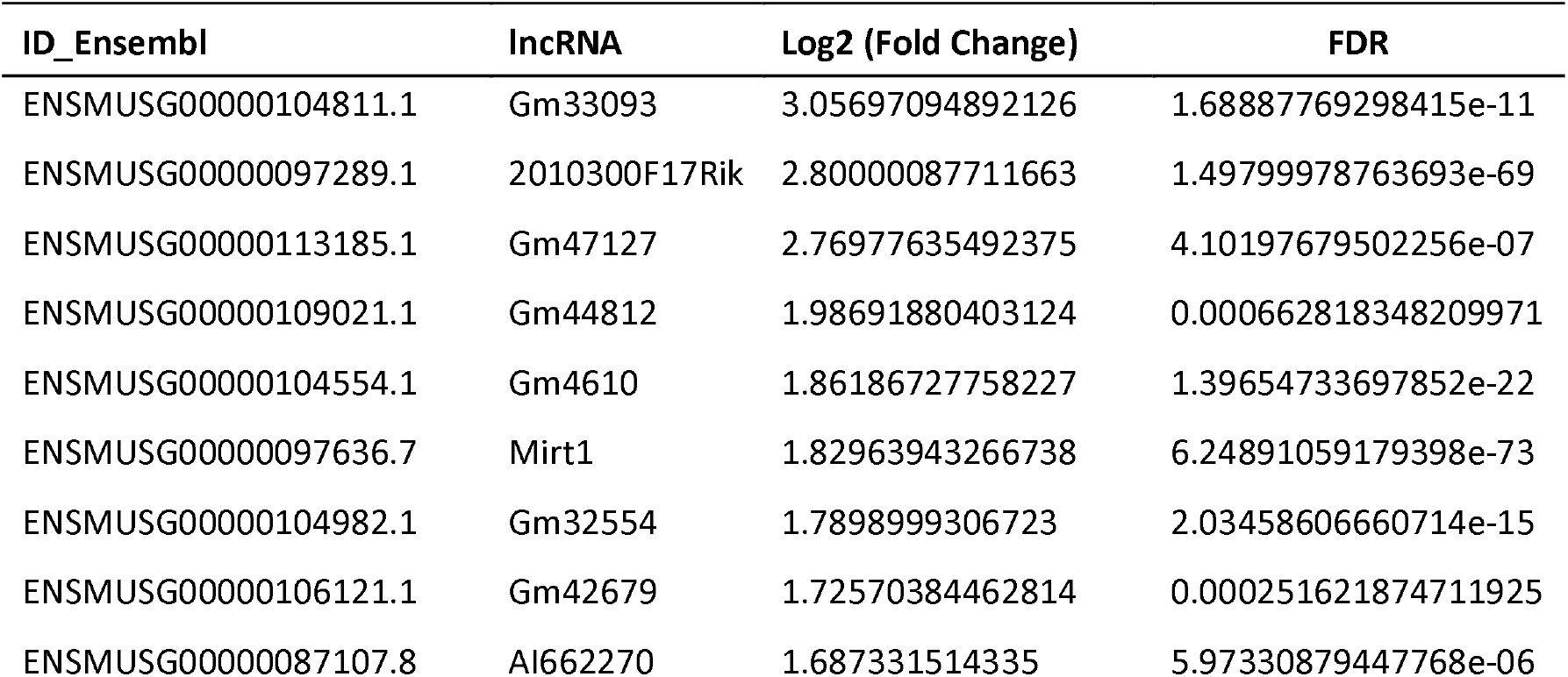
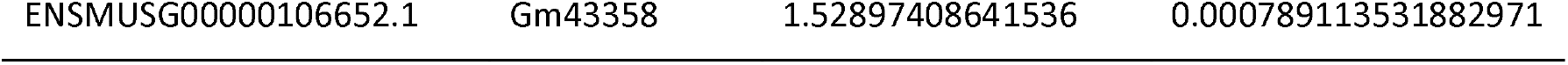
Top 10 upregulated differentially expressed lncRNAs in Aire KO mTECs.

**Supplementary Table 1.1.**
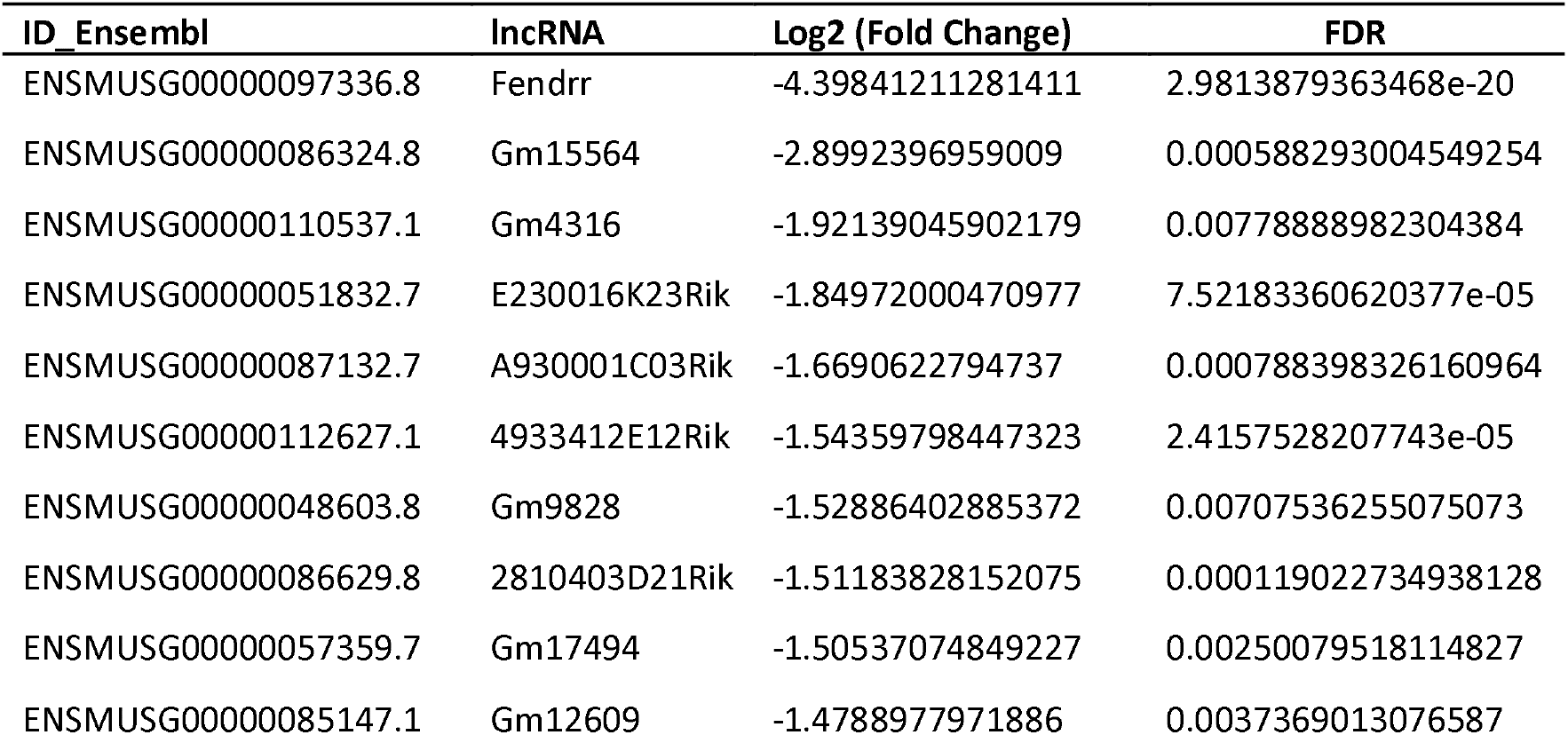
Top 10 downregulated differentially expressed lncRNAs in Aire KO mTECs.

**Supplementary Table 2.**
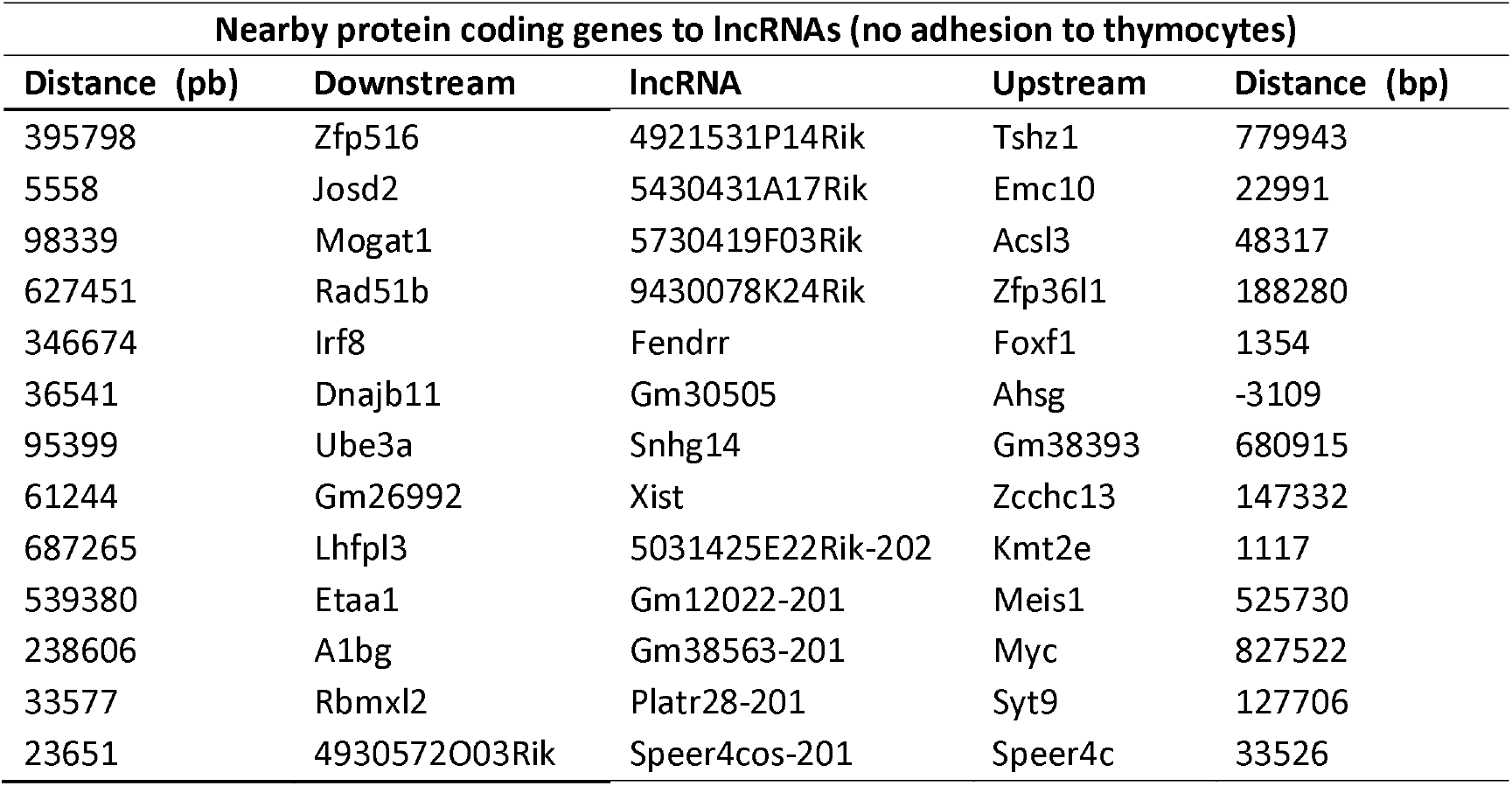
Nearby protein coding genes to the differentially expressed lncRNAs.

**Supplementary Table 2.1.**
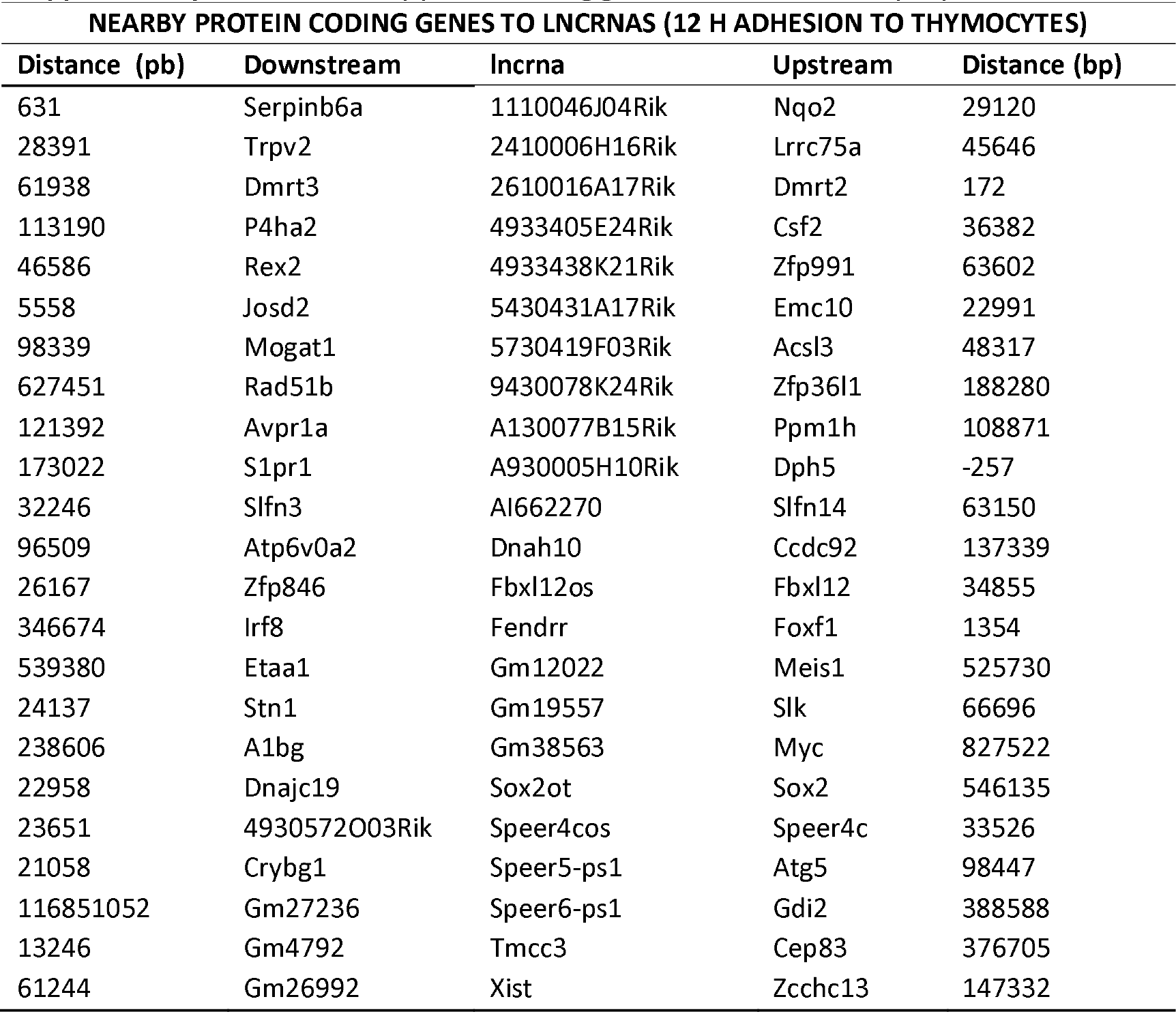
Nearby protein coding genes to the differentially expressed IncRNAs.

**Supplementary Table 2.2.**
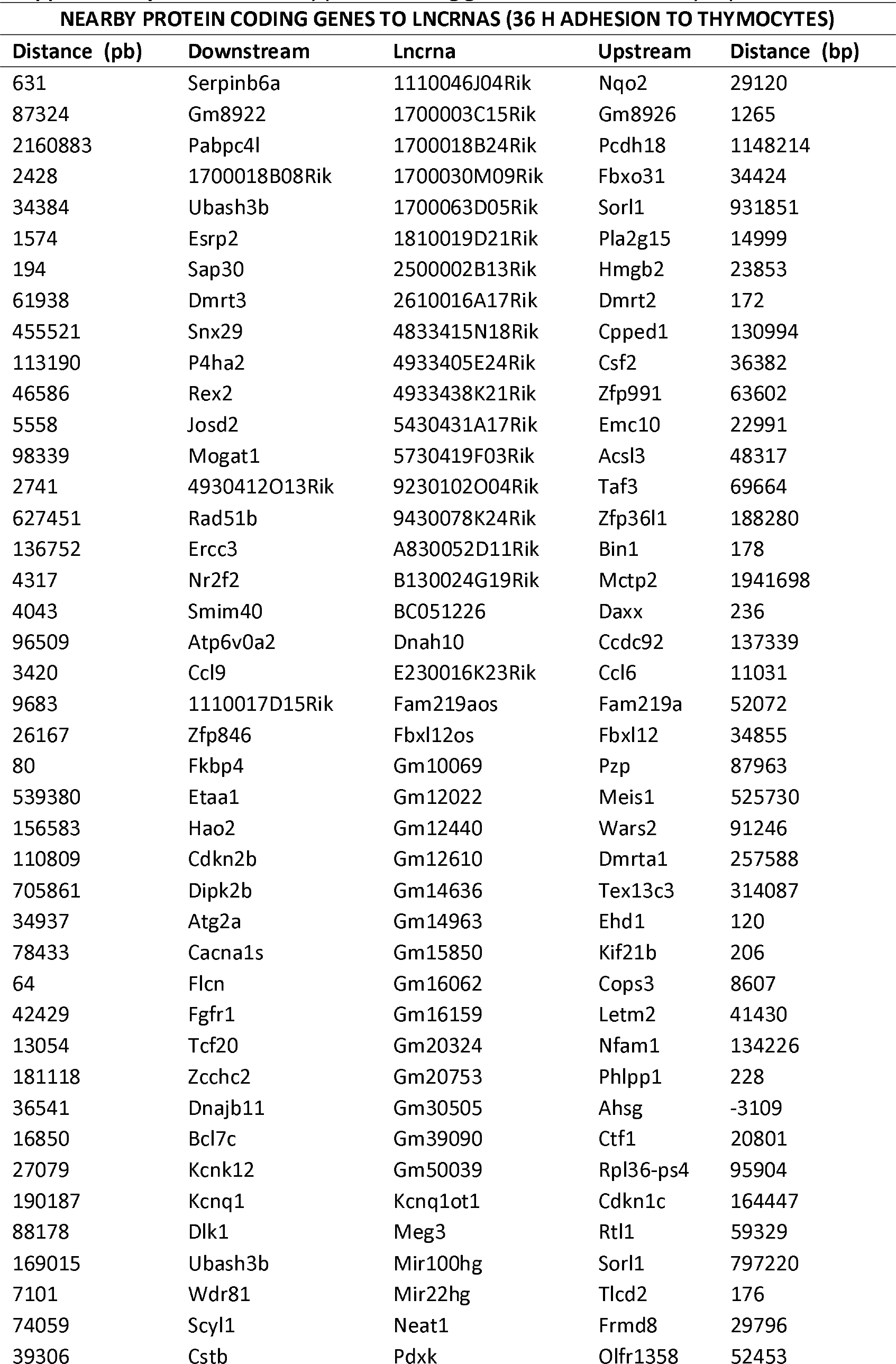
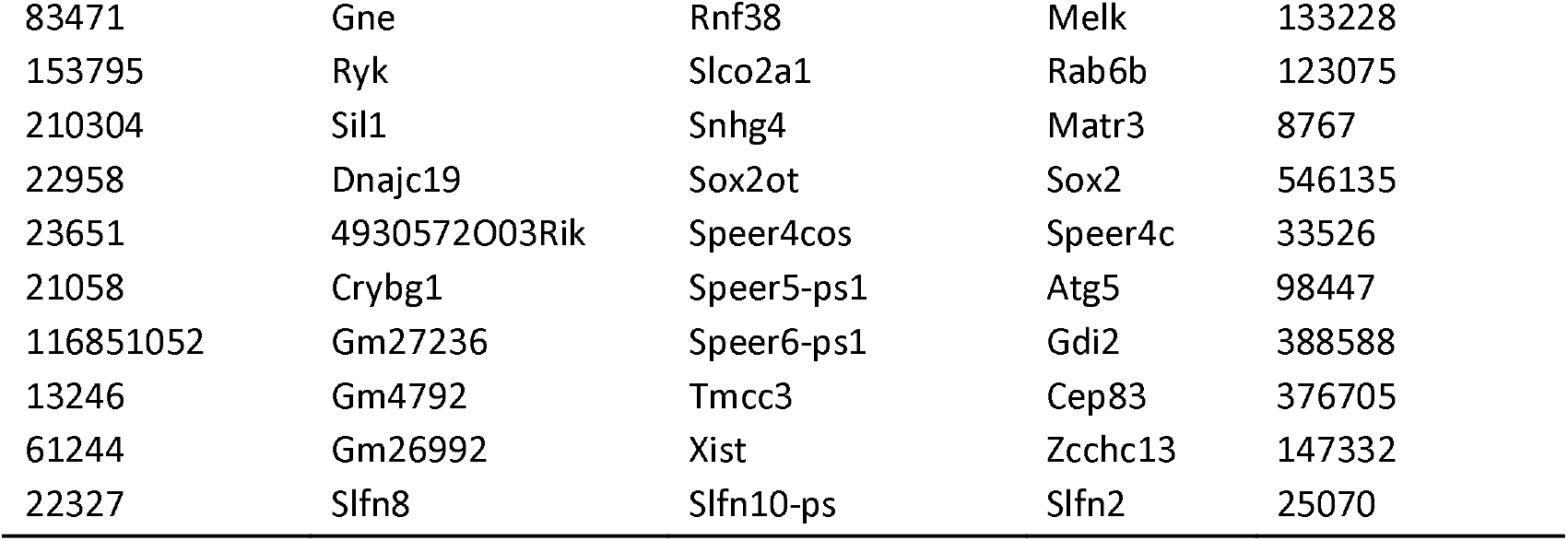
Nearby protein coding genes to the differentially expressed lncRNAs.

**Supplementary Table 3.** Coding potential of lncRNAs.

| lncRNA | Coding probability | Classification |
| --- | --- | --- |
| SNHG14 | 0.439997 | noncoding |
| FENDRR | 0.0656715 | noncoding |
| PLATR28 | 0.0305438 | noncoding |
| SPEER4COS-201 | 0.201759 | noncoding |
| 4921531P14RIK | 0.0398758 | noncoding |
| 5031425E22RIK | 0.424424 | noncoding |
| 5430431A17RIK | 0.212594 | noncoding |
| 5730419F03RIK | 0.030601 | noncoding |
| 9430078K24RIK | 0.157714 | noncoding |
| GM12022-201 | 0.0201895 | noncoding |
| GM30505 | 0.103945 | noncoding |
| GM38563-201 | 0.0475214 | noncoding |
| XIST | 0.0241659 | noncoding |

**Supplementary Table 3.1.**
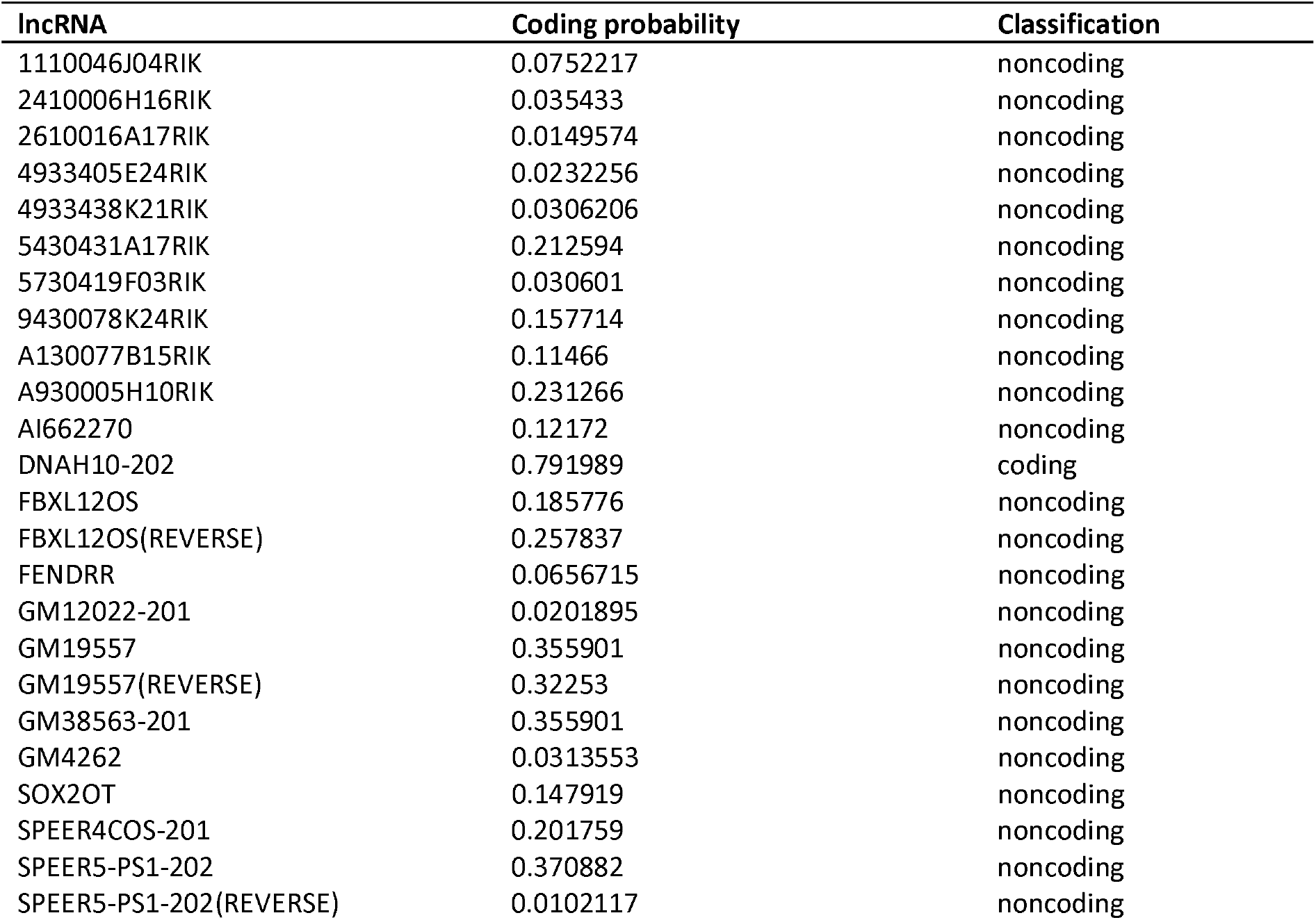
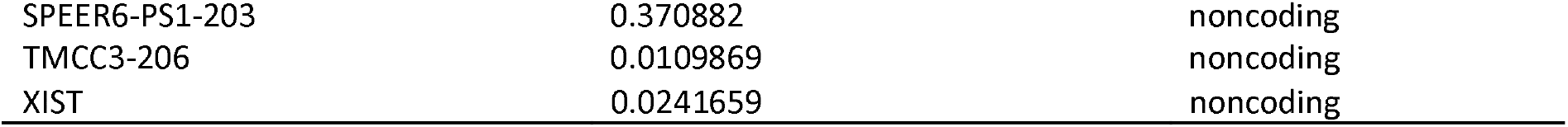
Coding potential of lncRNAs after 12h adhesion to thymocytes.

**Supplementary Table 3.2.** Coding potential of lncRNAs after 36h adhesion to thymocytes.

| lncRNA | Coding probability | Classification |
| --- | --- | --- |
| 1110046J04Rik | 0.0752217 | noncoding |
| 1700003C15Rik | 0.0418878 | noncoding |
| 1700018B24Rik | 0.595009 | coding |
| 1700030M09Rik | 0.0290032 | noncoding |
| 1700063D05Rik | 0.0000117516 | noncoding |
| 1810019D21Rik | 0.0222635 | noncoding |
| 2500002B13Rik | 0.101008 | noncoding |
| 2610016A17Rik | 0.0149574 | noncoding |
| 4833415N18Rik | 0.130809 | noncoding |
| 4930542C21Rik | 0.0281985 | noncoding |
| 4933405E24Rik | 0.0232256 | noncoding |
| 4933438K21Rik | 0.0701577 | noncoding |
| 5430431A17Rik | 0.212594 | noncoding |
| 5730419F03Rik | 0.030601 | noncoding |
| A830052D11Rik | 0.0141421 | noncoding |
| B130024G19Rik | 0.266976 | noncoding |
| BC051226 | 0.0939098 | noncoding |
| Dnah10-202 | 0.791989 | coding |
| E230016K23Rik | 0.64385 | coding |
| Fam219aos | 0.484752 | noncoding |
| Fbxl12os | 0.185776 | noncoding |
| Gm10069 | 0.645588 | coding |
| Gm10536 | 0.0343469 | noncoding |
| Gm12022-201 | 0.0201895 | noncoding |
| Gm12440 | 0.0262182 | noncoding |
| Gm12610 | 0.0263595 | noncoding |
| Gm14005_Morrbid | 0.156178 | noncoding |
| Gm14636 | 0.509347 | coding |
| Gm14963 | 0.0284942 | noncoding |
| Gm15850 | 0.0998467 | noncoding |
| Gm16062 | 0.586844 | coding |
| Gm16159 | 0.0252371 | noncoding |
| Gm20324 | 0.654366 | coding |
| Gm20753 | 0.195639 | noncoding |
| Gm30505 | 0.103945 | noncoding |
| Gm39090 | 0.088985 | noncoding |
| Gm4827 | 0.354224 | noncoding |
| Gm50039 | 0.776921 | coding |
| Kcnq1ot1 | 0.141033 | noncoding |
| Meg3 | 0.408619 | noncoding |
| Mir100hg | 0.0765838 | noncoding |
| Mir22hg | 0.0323014 | noncoding |
| Neat1 | 0.178003 | noncoding |
| Pdxk-ps-203 | 0.0433202 | noncoding |
| Rnf38-217 | 0.0000141858 | noncoding |
| Slco2a1-202 | 0.0717885 | noncoding |
| Slfn10-ps-201 | 0.093136 | noncoding |
| Snhg4 | 0.0246947 | noncoding |
| Sox2ot | 0.02483 | noncoding |
| Speer4cos-201 | 0.065561 | noncoding |
| Speer5-ps1-202 | 0.0124209 | noncoding |
| Speer6-ps1-203 | 0.0280487 | noncoding |
| Tmcc3-206 | 0.0109869 | noncoding |
| Xist_202 | 0.0241659 | noncoding |
| 9230102O04Rik-202 | 0.0244906 | noncoding |
| 9430078K24Rik | 0.106114 | noncoding |

